# Power and reproducibility in the external validation of brain-phenotype predictions

**DOI:** 10.1101/2023.10.25.563971

**Authors:** Matthew Rosenblatt, Link Tejavibulya, Chris C. Camp, Rongtao Jiang, Margaret L. Westwater, Stephanie Noble, Dustin Scheinost

## Abstract

Identifying reproducible and generalizable brain-phenotype associations is a central goal of neuroimaging. Consistent with this goal, prediction frameworks evaluate brain-phenotype models in unseen data. Most prediction studies train and evaluate a model in the same dataset. However, external validation, or the evaluation of a model in an external dataset, provides a better assessment of robustness and generalizability. Despite the promise of external validation and calls for its usage, the statistical power of such studies has yet to be investigated. In this work, we ran over 60 million simulations across several datasets, phenotypes, and sample sizes to better understand how the sizes of the training and external datasets affect statistical power. We found that prior external validation studies used sample sizes prone to low power, which may lead to false negatives and effect size inflation. Furthermore, increases in the external sample size led to increased simulated power directly following theoretical power curves, whereas changes in the training dataset size offset the simulated power curves. Finally, we compared the performance of a model within a dataset to the external performance. The within-dataset performance was typically within *r=0.2* of the cross-dataset performance, which could help decide how to power future external validation studies. Overall, our results illustrate the importance of considering the sample sizes of both the training and external datasets when performing external validation.

## 1. Introduction

Neuroimaging studies increasingly leverage large datasets to understand brain-phenotype associations (Horien *et al*., 2021). However, even traditionally “large” datasets, which include hundreds of participants, are underpowered for many association studies (Marek *et al*., 2022). Low statistical power presents numerous roadblocks to the reproducibility of neuroimaging research, including false negatives, inflated effect sizes, and replication failures (Yarkoni, 2009; Yarkoni and Braver, 2010; Button *et al*., 2013; Cremers, Wager and Yarkoni, 2017; Marek *et al*., 2022).

In contrast to association studies, prediction frameworks can alleviate the poor reproducibility seen in certain neuroimaging studies (Klapwijk *et al*., 2021; Rosenberg and Finn, 2022; Goltermann *et al*., 2023; Makowski *et al*., 2023; Spisak, Bingel and Wager, 2023). Unlike association, “prediction” entails the evaluation of a model on unseen data, which minimizes the risk of overfitting. Thus, it provides a more robust measure of brain-phenotype associations than in-sample associations. Typically, prediction is achieved by dividing a dataset into “training” and “test” sets, such as through k-fold cross-validation. Although an improvement over in-sample associations, splitting a dataset into training and test samples does not fully capture the generalizability and utility of brain-phenotype associations. Even with cross-validation, a model can be overfit to the idiosyncrasies of a particular dataset (Genon, Eickhoff and Kharabian, 2022; Yeung *et al*., 2022).

*External validation*, or applying a model to an entirely different dataset, is the gold standard when evaluating the generalizability of predictive models. Generalizing a model to another dataset with different characteristics provides strong evidence of a robust and reproducible brain-phenotype association. As such, numerous works encourage generalization to external datasets (Woo *et al*., 2017; Rosenberg, Casey and Holmes, 2018; Genon, Eickhoff and Kharabian, 2022; Rosenberg and Finn, 2022; Wu *et al*., 2022; Yeung *et al*., 2022). Since few studies have the resources to collect two independent samples, external validation is usually performed using an existing publicly available dataset. As the availability of such datasets continues to increase, external validation will likely become more accessible and commonplace.

Nevertheless, external datasets rarely harmonize with the primary dataset, often including differences in phenotypic measures or neuroimaging data. Researchers typically resort to the most similar dataset available. Given the limited number of options for external datasets, statistical power is rarely a consideration for external validation studies. Thus, the power of many external validation studies is unknown, and there remains a need for appropriate methodological approaches for determining the sample size required for external validation.

In this work, we explore how the sample sizes of both the training and external datasets affect cross-dataset prediction power in four large (n=424-7977), publicly available neuroimaging datasets. We first survey what training and external sample sizes have been used by existing external validation studies. Next, we resample the publicly available datasets across multiple sample sizes and evaluate internal (i.e., within-dataset) and external (i.e., across datasets) prediction performance. Finally, we investigate the relationship between the internal and external prediction performance.

## 2. Methods

### 2.1 Datasets

Resting-state fMRI data were obtained in each of our four datasets: the Adolescent Brain Cognitive Development (ABCD) Study (Casey *et al*., 2018), the Healthy Brain Network (HBN) Dataset (Alexander *et al*., 2017), the Human Connectome Project Development (HCPD) Dataset (Somerville *et al*., 2018), and the Philadelphia Neurodevelopmental Cohort (PNC) Dataset (Satterthwaite *et al*., 2014, 2016). Details about the datasets are presented in Table S1. In brief, the ABCD dataset consists of 9–10-year-olds who underwent fMRI scanning across 21 sites in the United States (n=7822-7977 across phenotypes). The HBN dataset consists of participants aged 5-22 years recruited from four sites near the New York greater metropolitan area (n=1024-1201). The HCPD dataset consists of participants aged 8-22 years who completed fMRI scanning across four sites in the United States (Harvard, UCLA, University of Minnesota, Washington University in St. Louis) (n=424-605). The PNC dataset consists of 8–21-year-olds in the Philadelphia area who received care at the Children’s Hospital of Philadelphia (n=1106-1126).

Throughout this work, we predicted age, attention problems, and matrix reasoning in these four datasets. These measures span a wide range of effect sizes, making them particularly useful for investigating power and effect size inflation. For the attention problems measure, we used the Child Behavior Checklist (CBCL) (Achenbach and Ruffle, 2000) Attention Problems Raw Score in ABCD, HBN, and HCPD. In PNC, we used the Structured Interview for Prodromal Symptoms (Miller *et al*., 2003): Trouble with Focus and Attention Severity Scale (SIP001, accession code: phv00194672.v2.p2). We used the WISC-V (Wechsler, 2014) Matrix Reasoning Total Raw Score in ABCD, HBN, and HCPD for the matrix reasoning measure. In PNC, we used the Penn Matrix Reasoning (Bilker et al., 2012; Moore et al., 2015) Total Raw Score (PMAT_CR, accession code: phv00194834.v2.p2). Summary statistics for these measures are presented in Table S1. While we used the Matrix Reasoning Raw Score in the main text, additional results using the Matrix Reasoning Scaled Score are presented in Table S2 and Figures S7-9.

### 2.2 Preprocessing

Data were pre-processed using BioImage Suite (Papademetris *et al*., 2006). This pre-processing included regression of covariates of no interest from the functional data, including linear and quadratic drifts, mean cerebrospinal fluid signal, mean white matter signal, and mean global signal. Additional motion control was applied by regressing a 24-parameter motion model—which included six rigid body motion parameters, six temporal derivatives, and the square of these terms—from the data. Subsequently, we applied temporal smoothing with a Gaussian filter (approximate cutoff frequency=0.12 Hz) and gray matter masking, as defined in common space (Holmes *et al*., 1998). Then, the Shen 268-node atlas (Shen *et al*., 2013) was applied to parcellate the denoised data into 268 nodes. Finally, we generated functional connectivity matrices by correlating each time series from pairs of nodes and applying the Fisher transform.

Data were excluded for poor data quality, missing nodes due to lack of full brain coverage, high motion (>0.2mm mean frame-wise displacement), or missing phenotypic data. After applying these exclusion criteria, 7977, 1201, 605, and 1126 participants remained in ABCD, HBN, HCPD, and PNC, respectively.

### 2.3 Data subsampling

For the within-dataset validation, the main dataset was resampled without replacement and split into two subsets: a group to train predictive models (training group) and a group to evaluate the performance of the predictive models (held-out group). We chose to evaluate within-dataset performance using a held-out group instead of k-fold cross-validation because the variability in k-fold performance approaches zero as the training sample size approaches the main dataset size. The held-out group size was 100 for HCPD, 200 for HBN and PNC, and 3000 for ABCD. The training group was randomly subsampled at various logarithmically spaced sample sizes (see Figure 2, Figure S4 for sample sizes). We resampled the main and external datasets for the cross-dataset validation. For each training sample, models were evaluated in random subsets of the external dataset of various sample sizes (see Figure 2, Figure S4 for sample sizes).

The resampling procedure was repeated 100 times for the main dataset, and the external dataset was resampled 100 times for each of these repeats. Thus, we performed 10,000 evaluations for each combination of the training dataset, external dataset, phenotype, training sample size, and external sample size. In total, this paper included over 60 million model evaluations. A summary of the resampling procedure is presented in Figure S1.

### 2.4 Regression models

We will refer to two types of results throughout this work: 1) within-dataset validation and 2) external validation. For within-dataset validation, we evaluated performance in a randomly selected held-out sample. Covariates (sex, motion, and age, if applicable) were first regressed from the training data. Then, a ridge regression model was trained using the top 1% of features most correlated with the outcome of interest (Pedregosa *et al*., 2011). Five-fold cross-validation was performed within the training set to select the L2 regularization parameter *α* (*α*=10^{-3,-2,-1,0,1,2,3}^). Afterward, the entire pipeline was applied to the held-out test data. Crucially, the covariate regression parameters and features obtained from the training set were applied to the test set to avoid data leakage (Snoek, Miletić and Scholte, 2019; Chyzhyk *et al*., 2022). For cross-dataset validation, we used the same models as above. However, the model was evaluated with the external dataset instead of the held-out test data. Performance was evaluated with Pearson’s correlation *r* as it is among the most common measures used in neuroimaging predictive studies. For instance, Yeung et al. found that 97 of the 108 investigated studies used Pearson’s correlation as the evaluation metric (Yeung *et al*., 2022).

We will define the “ground truth” prediction performance as follows. For within-dataset predictions, the ground truth refers to the performance in the total sample averaged over 100 random iterations of nested 5-fold cross-validation. The ground truth was operationalized for external predictions as the prediction performance when training in the whole primary dataset and testing with the entire external dataset.

### 2.5 Predictive power calculation

We calculated predictive power for all combinations of training dataset, test dataset, and phenotype that had a significant ground truth effect. Since external validation involves testing a model in an independent dataset, directly converting *r* to *p-*values is appropriate, as opposed to cross-validation, where calculating *p*-values requires permutation testing. One-tailed significance testing was used since we only hypothesize that *r*>0 to achieve significant prediction performance. To calculate power in cross-dataset predictions, we computed the fraction of subsamples that achieved a significant prediction performance, as defined by the field-wide practice of *p*<0.05.

Furthermore, we compared the simulated power to the “theoretical power,” which assumes that the ground truth effect size is known. The theoretical power curve was calculated as:

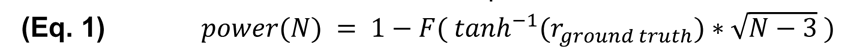

where *F* is the standard normal cumulative distribution function, *r_ground truth_* is the ground truth cross-dataset prediction performance, which we defined as the cross-dataset prediction performance using the full training and test datasets, and *N* is the sample size.

### 2.6 False positive rate

We computed the false positive rate for all cross-dataset predictions that did not have a significant ground truth effect. The false positive rate is the proportion of simulated examples for which the observed effect is significant (*p*<0.05) despite a ground truth effect that is not significant.

### 2.7 Performance effect size inflation

Another important consideration is the inflation of reported effect sizes, as documented by numerous previous studies (Yarkoni, 2009; Button *et al*., 2013; Cremers, Wager and Yarkoni, 2017; Marek *et al*., 2022). Low power reduces the likelihood of detecting an actual effect and leads to the inflation of reported significant effects (Yarkoni, 2009; Button *et al*., 2013). In other words, if significant results are reported in a low-powered sample, such as due to a small sample size, then the effect size is likely inflated.

We first examined all results that achieved significant prediction performance to approximate the inflation of effect sizes because this aligns with publication bias surrounding positive results. We agree with other works that non-significant results should still be published (Dwan *et al*., 2008; Button *et al*., 2013), but the current reality of the field is that most published results are significant predictions. Among the significant prediction results, we compared the effect size to the ground truth effect size and calculated the inflation relative to the ground truth (*Δr=r_reported_ -r_ground truth_*).

### 2.8 Relating internal and external performance

After looking at within-dataset performance and cross-dataset performance separately, we compared the two to determine whether within-dataset performance could inform how well a model would generalize. We calculated the difference between the within-dataset held-out performance (*r_internal_*) and the performance in the full external dataset (*r_external_*) for each training sample. We then assessed the performance difference across 100 iterations of random subsampling for each training dataset size.

### 2.9 Literature review of external validation sample sizes

We performed a brief literature review of sample sizes in neuroimaging external validation studies published in 2022-2023 to investigate the simulated power at typical sample sizes in the field. Supplemental Information Section *S5: Literature review of external validation sample sizes* provides the details of this review.

## 3. Results

In the main text, we show the results of training in the HBN dataset and testing in other datasets. All possible combinations of training/test datasets are included in the supplemental information.

### 3.1 External validation sample sizes in the literature

Among 27 qualifying articles published in 2022-2023, the median sample size of the training dataset was n=161 (IQR: 100-495), and the median sample size of the external dataset was n=94 (IQR: 39.5-682). A previous analysis by Yeung et al. included papers before 2022 (Yeung *et al*., 2022), finding 27 articles using external validation. In this sample, the median sample size of the training dataset was n=87 (IQR: 25-343), and the median sample size of the external dataset was n=137 (IQR: 60-197). Across both samples, the median training sample size was n=129 (IQR: 59.5-371.25), and the median external sample size was n=108 (IQR: 50-281).

### 3.2 Within-dataset performance

As the training sample size increased, within-dataset prediction performance also increased on average (representative HBN results in Figure 1; additional results in Figure S2). Unsurprisingly, variability in performance was greater at small sample sizes across all datasets and phenotypes (Figure S2).

**Figure 1.**
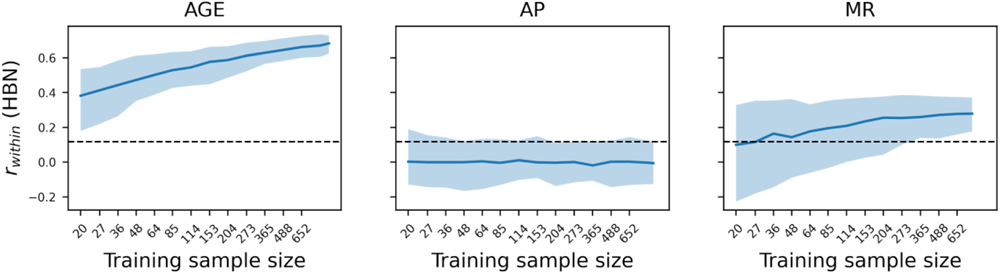
Within-dataset held-out prediction performance in HBN for age, attention problems, and matrix reasoning. The performance was evaluated in a randomly selected held-out sample of size n=200. The error bars show the 2.5^th^ and 97.5^th^ percentiles among 100 repeats of resampling at each training sample size. The dotted line reflects the correlation value required for a significance level of *p*<0.05. Similar results were observed for the ABCD, HCPD, and PNC datasets; see Figures S2-3. AP: attention problems, MR: matrix reasoning.

Because of this variability, effect sizes at small sample sizes were sometimes greater than the ground truth. For example, at a sample size of n=204 in HBN, the fraction of subsamples with prediction performance of *Δr*>0.05 compared to the ground truth was 0% for age, 11% for attention problems, and 24% for matrix reasoning. Similar trends were seen across all datasets (Figure S2), where the highest proportion of effect size inflation occurred in attention problems and matrix reasoning prediction. Still, there was little to no inflation for age prediction. Furthermore, the inflation of effects was rare in ABCD, which had by far the largest held-out group.

### 3.3 Baseline cross-dataset performance

Along with within-dataset performance, we evaluated cross-dataset performance. Ground truth performances for each dataset and phenotype—evaluated using the full training and external dataset sizes—varied from non-predictive to strong (Table 1). All age models significantly predicted across datasets, and all matrix reasoning models cross-predicted, except for when testing in ABCD. Three of the twelve attention problems models had weakly significant performance. Notably, we evaluated the cross-dataset performance even when the within-dataset performance was not significant for the sake of completeness.

**Table 1.** Ground truth performance for cross-dataset predictions using full training and external samples. AP: attention problems, MR: matrix reasoning. *p<0.05, **p<1e-5.

|  | Training Data |  |  |  |  |  |  |  |  |  |  |  |
| --- | --- | --- | --- | --- | --- | --- | --- | --- | --- | --- | --- | --- |
|  | ABCD |  |  | HBN |  |  | HCPD |  |  | PNC |  |  |
| External Data | Age | AP | MR | Age | AP | MR | Age | AP | MR | Age | AP | MR |
| ABCD | Within | Within | Within | N/A | 0.02 | -0.03 | N/A | 0.02 | 0.03* | N/A | 0.04* | 0.03* |
| HBN | N/A | -0.01 | 0.29** | Within | Within | Within | 0.58** | 0.00 | 0.31** | 0.54** | -0.02 | 0.26** |
| HCPD | N/A | 0.07 | 0.43** | 0.73** | 0.05 | 0.25** | Within | Within | Within | 0.65** | 0.00 | 0.26** |
| PNC | N/A | 0.09* | 0.23** | 0.48** | 0.00 | 0.22** | 0.42** | 0.06* | 0.20** | Within | Within | Within |

### 3.4 Power and false positive rate for cross-dataset predictions

In all datasets, cross-dataset prediction power was affected by both the external dataset size and the training dataset size (representative HBN results in Figure 2; additional results in Figure S4). Furthermore, when assuming the ground truth effect size was known, the cross-dataset power followed the theoretical curve for power of correlations (Figure 2; Figure S4; see blue lines). Decreasing the size of the training dataset appeared to negatively offset the theoretical power curve.

**Figure 2.**
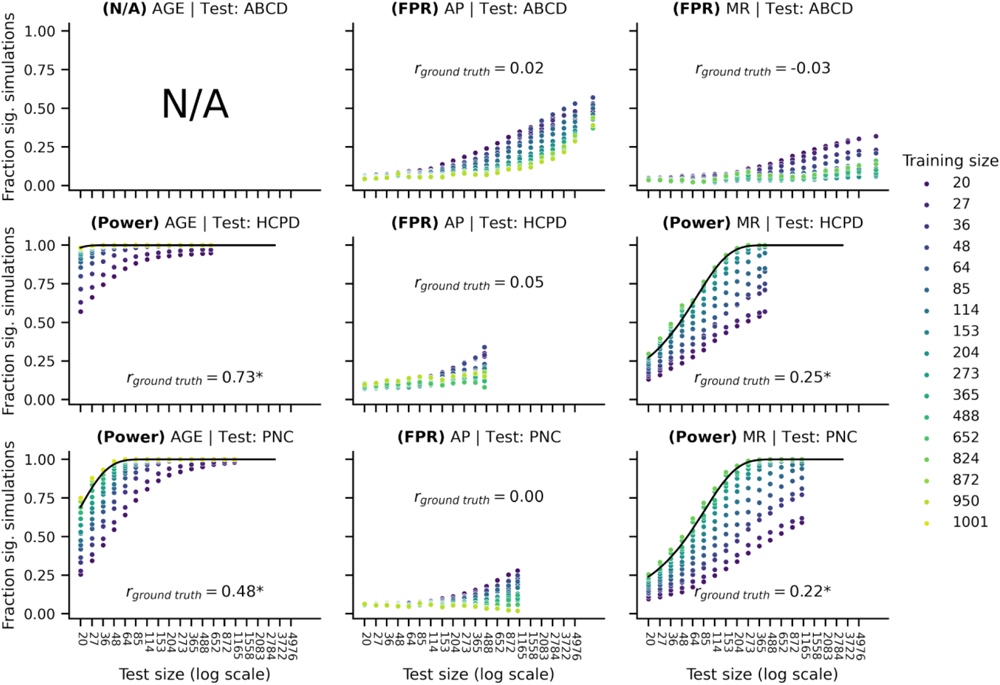
Power and false positive rates for cross-dataset predictions, training in HBN and testing in ABCD (top row), HCPD (middle row), or PNC (bottom row) for prediction of age (left column), attention problems (middle column), or matrix reasoning (right column). The blue lines represent theoretical power assuming a known ground truth performance. The panel with N/A means that data were not included in this study. Similar results were observed for the ABCD, HCPD, and PNC datasets; see Figure S4. AP: attention problems, MR: matrix reasoning.

For cases where the ground truth effect was non-significant, we found that the false positive rate was highest for large external samples and small training samples. At large sample sizes, effects can achieve significance with a very small effect size. Thus, with the high variability of training samples at a small sample size, there is a risk of fitting a “lucky” model, leading to false positives.

Across all datasets, age had the highest ground truth effect size (*r*=0.42-0.73). It could achieve more power with fewer test samples than attention problems or matrix reasoning, which directly follows Equation 1. Furthermore, greater power was achieved with smaller training samples in age predictions relative to attention problems or matrix reasoning. This result suggests that strong effects, such as age, can be robustly detected in small samples. Notably, using the full external samples but training samples of only n=20, all six cross-dataset age predictions had power ranging from 86-100%. However, as described above, small training and large test samples pose the greatest risk for false positives in cases where the effect size is smaller.

We also tested power for the median sample sizes based on our literature review. The training sample size closest to the median was n=114 and the external sample size closest to the median was n=114. For these sample sizes, the power ranged across training/external dataset combinations from 99.11-100.00% for age, 5.47-8.35% for attention problems, and 5.24-72.74% for matrix reasoning. For sample sizes comparable to the 25^th^ percentile in the field (training size: n=64, test size: n=48), the power was 78.33-98.94% for all dataset combinations for age, 4.86-6.84% for attention problems, and 5.67-35.63% for matrix reasoning. When instead considering sample sizes comparable to the 75^th^ percentile in the field (training size: n=365, test size: n=273), the power was 100.00% for all dataset combinations for age, 8.34-9.50% for attention problems, and 8.22-99.57% for matrix reasoning. In particular for attention problems and matrix reasoning, common sample sizes for external validation in the field appear to be underpowered, where 80% power is the typical goal.

### 3.5 Effect size inflation for cross-dataset predictions

Among significant results, we computed the median effect size inflation (or deflation) relative to the ground truth (representative HBN results in Figure 3; additional results in Figure S5). Across all datasets, effect size inflation was greatest in weaker predictions and smallest in strong predictions, such as age. For the weakest predictive models, the training dataset size made little difference in effect size inflation, likely because effect size inflation is a consequence of low power based on the *test sample size*. For stronger models (e.g., age), we saw a greater effect of training size. There was little to no inflation, but smaller training sizes produced worse predictions. When predicting age, we previously mentioned that >80% power could be achieved with small training samples and large external samples. Still, the deflation of effects shows the primary disadvantage of using small training samples.

**Figure 3.**
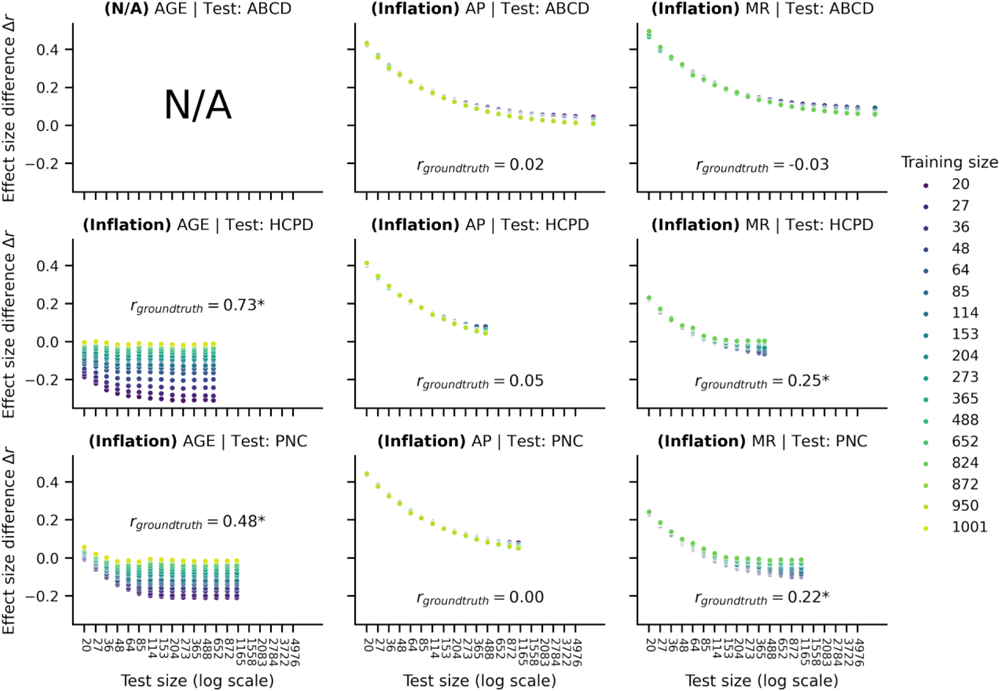
Median effect size inflation for cross-dataset predictions, training in HBN and testing in ABCD (top row), HCPD (middle row), or PNC (bottom row) for prediction of age (left column), attention (middle column), or matrix reasoning (right column). Panels with N/A mean that data were not available. Similar results were observed for the ABCD, HCPD, and PNC datasets; see Figure S5. AP: attention problems, MR: matrix reasoning.

Using the training sample size closest to the median in the field (n=114), the external sample size closest to the field-wide median (n=114) showed median inflation rates, where negative inflation means deflation, ranging across datasets from *Δr* of -0.12 to -0.05 for age, 0.10 to 0.20 for attention problems, and -0.17 to 0.21 for matrix reasoning. If using smaller external sample sizes, such as that closest to the 25^th^ percentile in the field (training size: n=64, test size: n=48), the inflation rates ranged from -0.16 to -0.05 for age, 0.20 to 0.31 for attention problems, and -0.10 to 0.32 for matrix reasoning. For sample sizes comparable to the 75^th^ percentile (training size: n=365, test size: n=273), the inflation rates were -0.06-0.00 for age, 0.03-0.14 for attention problems, and -0.17-0.15 for matrix reasoning. For age and similar strong predictions, typical sample sizes in the field could lead to underestimating effect sizes. In contrast, effect sizes may be overestimated for attention problems and matrix reasoning.

### 3.6 Relating within-to cross-dataset performance

A key remaining question is how within-dataset and cross-dataset performance may be related, and whether a possible association can inform future cross-dataset studies. As such, we compared the within-dataset held-out performance (*r_internal_*) to the performance in the full external dataset (*r_external_*) for each training subsample (representative HBN results in Figure 4; additional results in Figure S6). In most cases, the average within-dataset performance was within *r=0.2* of the cross-dataset prediction. Although the average was a relatively good estimate of the cross-dataset performance, we do not have the luxury of averaging across many different subsamples in neuroimaging. The difference in internal and external performances was highly variable for any given subsample, especially at smaller sample sizes.

**Figure 4.**
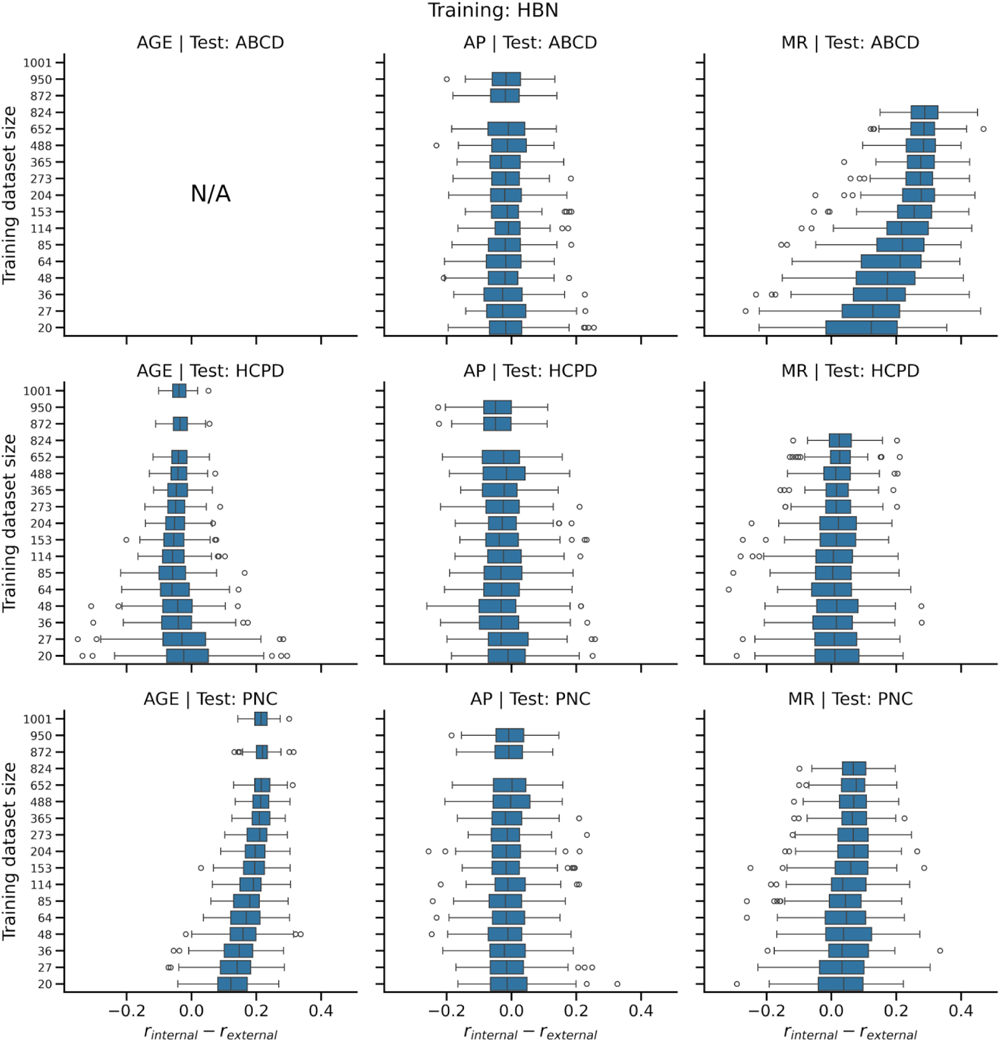
Boxplots of the difference between internal and external performance for each subsample of the training data. For each training data size, 100 random subsamples were taken. The model was evaluated for internal performance in a held-out sample of size n=200. For external performance, the model formed in the training subsample was applied to the full external dataset. Panels with N/A mean that data were not available. Similar results were observed for the ABCD, HCPD, and PNC datasets; see Figure S6. AP: attention problems, MR: matrix reasoning.

The internal and external performance were not always closely related on average. In particular, matrix reasoning predictions did not generalize to ABCD, so *r_internal_* - *r_external_* was consistently greater than zero. Inversely, matrix reasoning models from ABCD generalized to the other datasets more strongly than the within-dataset performance, so *r_internal_* - *r_external_* was negative.

When deciding how to power an external validation study, one should most heavily consider cases where *r_internal_* is much greater than *r_external_*, which would lead to false negatives or potential effect size inflation. At the training size closest to the existing median in the field (n=114), 86.57% of evaluations across all datasets and phenotypes met the requirement of (*r_internal_* - *r_external_* < 0.2), and 71.10% met the criteria when restricting to (*r_internal_* - *r_external_* < 0.1). At the sample size closest to the 25^th^ percentile of existing studies (n=64), 88.23% of studies were within the threshold of 0.2, and 72.57% were within the threshold of 0.1. At the sample size closest to the 75^th^ percentile of existing studies (n=365), 83.42% and 71.83% were within the thresholds of 0.2 and 0.1, respectively. Counterintuitively, using more training data resulted in internal prediction performance that was less consistent with the external performance for each subsample. This trend is partially due to smaller sample sizes having worse average internal *and* external performance. As such, if the data are restricted to results that only obtain within-dataset significance, the ratio of internal to external performance *r_internal_* / *r_external_* was less than 1.2 in 53.45% of evaluations for n=64, 53.81% for n=114, and 57.52% for n=365. The ratio was less than 1.5 in 67.80%, 68.83%, and 74.95% of evaluation for n=64, 114, and 365, respectively. Smaller samples tend to have the largest fractional increase in internal relative to external performance with increasing training sample size, suggesting that internal performance may be especially inflated relative to external performance when using small sample sizes.

## 4. Discussion

This work investigated power and effect size inflation in predictive models of brain-phenotypic associations as a function of training and external dataset sizes. Our results suggest that prior external validation studies have relied on sample sizes prone to low power, potentially leading to false negatives and effect size inflation. Increasing the sample size of external datasets increased the power following theoretical curves, whereas the training dataset size offset the power curve. Relatedly, false positive findings were most frequent for non-significant ground truth effects when using small training and large external datasets. For attention problems and matrix reasoning, significant effects were inflated with smaller external dataset sizes. However, for age, which exhibited the largest effect size, there was deflation when using small training samples. Finally, the within-dataset performance was usually within *r=0.2* of the cross-dataset performance. These results serve two purposes. First, they contextualize existing external validation results in the predictive neuroimaging literature. Second, they underscore potential pitfalls when implementing external validation in future studies.

Though external validation only occurs in a minority of neuroimaging prediction studies (Yeung *et al*., 2022), we expect that it will become increasingly prominent as the field confronts ongoing reproducibility challenges. In addition, external validation may help to ameliorate machine learning ethical issues (Mitchell *et al*., 2019; Chandler, Foltz and Elvevåg, 2020), including bias (Benkarim *et al*., 2021; Greene *et al*., 2022; Li *et al*., 2022) and trustworthiness (Rosenblatt *et al*., 2023). For bias, evaluating models in external datasets will better depict the robustness and generalizability of brain-phenotype associations in populations with different characteristics (Mehrabi *et al*., 2021; Tejavibulya *et al*., 2022). For trustworthiness, external validation ensures that data manipulations are not driving the results (Finlayson *et al*., 2019; Rosenblatt *et al*., 2023). Given the promise of external validation for improving reproducibility, bias, and trustworthiness, neuroimaging may follow a similar trajectory as genome-wide association studies, for which external replication is now a standard practice (Poldrack *et al*., 2017; Uffelmann *et al*., 2021).

Adequately powered studies mitigate against potential false negatives and effect size inflation, which, in turn, promotes the reproducibility and utility of scientific insights (Yarkoni, 2009; Yarkoni and Braver, 2010; Button *et al*., 2013; Cremers, Wager and Yarkoni, 2017; Marek *et al*., 2022). While large training datasets are needed to avoid overfitting or poor generalizability, the external dataset sample size is arguably more important for power in cross-dataset predictions. The power is proportional to the square root of the external sample size, but it only indirectly depends on the training sample size via the quality of the model. Furthermore, smaller training datasets are applicable when the brain-phenotype associations are strong. As such, reproducible brain-phenotype associations require large sample or effect sizes (Gratton, Nelson and Gordon, 2022). As an extreme example, age predictions with a training size of only n=20 had power ranging from 86-100% when using the full external dataset. Still, we would not recommend using a small training sample in cross-sectional external validation studies. The combination of small training samples (<100) and large external samples (>500) increased the likelihood of false positives.

In addition to power, effect size—measured by correlation—is another crucial component of external validation. Intuitively, smaller external dataset sizes require larger effect sizes to achieve significance. Combined with the reporting bias toward significant effects (Greenwald, 1975; Munafò, Stothart and Flint, 2009; Button *et al*., 2013; Open Science Collaboration, 2015), published effects with small test or external datasets may be inflated. Encouraging researchers to publish the results of external validation attempts—regardless of statistical significance— would ameliorate this issue. However, a more realistic solution could be to promote the use of large external dataset sizes. Effect sizes are unlikely to be inflated in large external test sets. One caveat is that statistical significance can be achieved with trivial effect sizes. For instance, a significant effect of *r*=0.03, n=5000 may not be very meaningful, but it has a p-value less than 0.05. However, it is not to say that small effects cannot be meaningful, as these can affect policy (Searle *et al*., 2014; Gratton, Nelson and Gordon, 2022) or inform our understanding of a more complex characteristic. Instead, we emphasize that reporting and interpreting the effect size and significance are crucial in understanding brain-phenotype associations in large datasets (Cohen, 1994; Gigerenzer, 2004).

If the ground truth effect size for a given cross-dataset brain-phenotype association was known, the required sample size could be calculated directly using power curves. Unfortunately, perfect knowledge of the ground truth effect size would require evaluating the cross-dataset prediction before the study. Instead, one must rely on either within-dataset prediction performance (if the main dataset has already been collected) or published effect sizes, which typically represent within-dataset prediction rather than external validation. Based on our results, accounting for the drop-off in external dataset predictions by subtracting 0.1 to 0.2 from the within-dataset or literature correlation values may be a quick and dirty rule of thumb. A decrease in external validation prediction performance compared to within-dataset prediction is generally expected due to dataset shift, which is when the training and test populations are mismatched in a way that may degrade performance (Subbaswamy and Saria, 2020; Dockès, Varoquaux and Poline, 2021; Finlayson *et al*., 2021). A mismatch between datasets may come from differences in population characteristics, image acquisition, or phenotypic measurements. If the training and external datasets are too dissimilar, a rule of thumb might not account for dataset shift.

There were several limitations to our study. First, we focused on external validation instead of replication in an independent sample. Whereas external validation involves applying a model to another dataset, replication in an independent sample entails repeating the entire analysis in an independent dataset. Both are valid strategies to improve reproducibility and replicability, but from a predictive sense, external validation is more common. Second, we only analyzed multivariate brain-phenotype associations, as multivariate patterns are more reliable and becoming more popular than univariate associations. Third, to evaluate within-dataset performance, we used a small held-out sample (as small as n=100). This limitation was due to the size of the datasets, but we repeated the evaluation for 100 different random subsamples of size n=100 to reduce the noise. Fourth, the datasets in our study are all relatively similar. All participants live in the United States, are youths, and were born to the same generation. There are still differences between these datasets—the region within the United States, clinical diagnosis, and specific measurements. Whether our results generalize to datasets with other differences remains to be seen. Fifth, we studied the external validation of cross-sectional brain-phenotype associations. Still, other studies, such as longitudinal ones, may have greater power with smaller sample sizes (Gratton, Nelson and Gordon, 2022).

When selecting a dataset for external validation of a predictive model, one may have few options, depending on the phenotype of interest. If one must use a small training or external dataset in an external validation study, recognizing and explicitly acknowledging the sample size limitations will be crucial for promoting reproducibility. Despite the current reliance of the field on within-dataset associations and predictions, external validation will become more widespread. This work provides a starting point for understanding what sample sizes are required to power external validation studies adequately.

## Data and code availability

Data are available through the Adolescent Brain Cognitive Development Study (Casey *et al*., 2018), the Healthy Brain Network Dataset (Alexander *et al*., 2017), the Human Connectome Project Development Dataset (Somerville *et al*., 2018), and the Philadelphia Neurodevelopmental Cohort Dataset (Satterthwaite *et al*., 2014, 2016). Code for the analyses is available at: https://github.com/mattrosenblatt7/external_validation_power.

## Acknowledgements

This study was supported by the National Institute of Mental Health grant R01MH121095 (obtained by D.S.). M.R. was supported by the National Science Foundation Graduate Research Fellowship under grant DGE2139841. L.T. was supported by the Gruber Science Fellowship. S.N. was supported by the National Institute of Mental Health under grant K00MH122372. Any opinions, findings, and conclusions or recommendations expressed in this material are those of the authors and do not necessarily reflect those of the funding agencies.

The Human Connectome Project Development data was supported by the National Institute Of Mental Health of the National Institutes of Health under Award Number U01MH109589 and by funds provided by the McDonnell Center for Systems Neuroscience at Washington University in St. Louis. The HCP-Development 2.0 Release data used in this report came from DOI: 10.15154/1520708. Additional data used in the preparation of this article were obtained from the Adolescent Brain Cognitive Development (ABCD) Study (https://abcdstudy.org), held in the NIMH Data Archive (NDA). This is a multisite, longitudinal study designed to recruit more than 10,000 children age 9-10 and follow them over 10 years into early adulthood. The ABCD Study® is supported by the National Institutes of Health and additional federal partners under award numbers U01DA041048, U01DA050989, U01DA051016, U01DA041022, U01DA051018, U01DA051037, U01DA050987, U01DA041174, U01DA041106, U01DA041117, U01DA041028, U01DA041134, U01DA050988, U01DA051039, U01DA041156, U01DA041025, U01DA041120, U01DA051038, U01DA041148, U01DA041093, U01DA041089, U24DA041123, U24DA041147.

A full list of supporters is available at https://abcdstudy.org/federal-partners.html. A listing of participating sites and a complete listing of the study investigators can be found at https://abcdstudy.org/consortium_members/. ABCD consortium investigators designed and implemented the study and/or provided data but did not necessarily participate in the analysis or writing of this report. This manuscript reflects the views of the authors and may not reflect the opinions or views of the NIH or ABCD consortium investigators. The Healthy Brain Network (http://www.healthybrainnetwork.org) and its initiatives are supported by philanthropic contributions from the following individuals, foundations and organizations: Margaret Bilotti; Brooklyn Nets; Agapi and Bruce Burkard; James Chang; Phyllis Green and Randolph Cōwen; Grieve Family Fund; Susan Miller and Byron Grote; Sarah and Geoff Gund; George Hall; Jonathan M. Harris Family Foundation; Joseph P. Healey; The Hearst Foundations; Eve and Ross Jaffe; Howard & Irene Levine Family Foundation; Rachael and Marshall Levine; George and Nitzia Logothetis; Christine and Richard Mack; Julie Minskoff; Valerie Mnuchin; Morgan Stanley Foundation; Amy and John Phelan; Roberts Family Foundation; Jim and Linda Robinson Foundation, Inc.; The Schaps Family; Zibby Schwarzman; Abigail Pogrebin and David Shapiro; Stavros Niarchos Foundation; Preethi Krishna and Ram Sundaram; Amy and John Weinberg; Donors to the 2013 Child Advocacy Award Dinner Auction; Donors to the 2012 Brant Art Auction. Additional data were provided by the PNC (principal investigators Hakon Hakonarson and Raquel Gur; phs000607.v1.p1). Support for the collection of these datasets was provided by grant RC2MH089983 awarded to Raquel Gur and RC2MH089924 awarded to Hakon Hakonarson.

## Supplemental Information

### S1. Dataset summaries

**Table S1.** Summary of the four datasets and three phenotypes used in this work. The proportions of male/female participants reflect self-reported sex. AP: attention problems; MR: matrix reasoning.

|  | ABCD (n=7977;<br>49.17% female) |  |  | HBN (n=1201;<br>39.80% female) |  |  | HCPD (n=605;<br>53.72% female) |  |  | PNC (n=1126;<br>54.62% female) |  |  |
| --- | --- | --- | --- | --- | --- | --- | --- | --- | --- | --- | --- | --- |
|  | Age | AP | MR | Age | AP | MR | Age | AP | MR | Age | AP | MR |
| Mean | 9.92 | 2.91 | 18.05 | 11.65 | 7.41 | 18.36 | 14.61 | 2.03 | 21.08 | 14.80 | 1.03 | 11.99 |
| SD | 0.62 | 3.46 | 3.76 | 3.42 | 4.54 | 4.46 | 3.90 | 2.56 | 3.96 | 3.29 | 1.19 | 4.09 |
| Range | 9.00-10.92 | 0.00-20.00 | 0.00-30.00 | 5.00-22.00 | 0.00-19.00 | 2.00-31.00 | 8.08-21.92 | 0.00-18.00 | 11.00-31.00 | 8.00-21.00 | 0.00-6.00 | 0.00-24.00 |
| # Available | 7977 | 7976 | 7822 | 1201 | 1150 | 1024 | 605 | 462 | 424 | 1126 | 1106 | 1119 |

### S2. Sampling procedure

**Figure S1.**
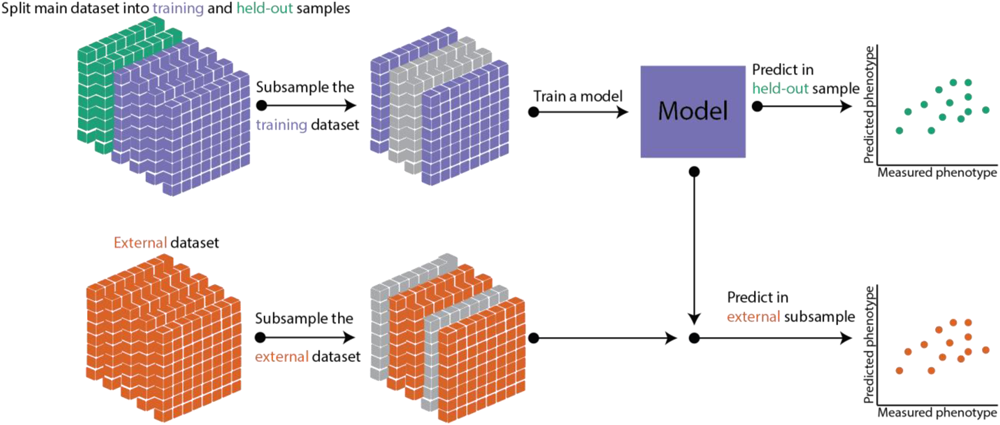
Summary of subsampling procedure in external validation. The main dataset was first split into two subsets: a group to train predictive models (training group) and an evaluation group (held-out group). We then subsampled the training dataset at various sample sizes and trained a model. The model was evaluated in the held-out group to estimate within-dataset performance. An external dataset was also subsampled at various sample sizes. The model was evaluated in these external subsamples to estimate external validation performance. The subsampling procedure was repeated 100 times for the main dataset, and the external dataset was subsampled 100 times for each of these repeats. Thus, we performed 10,000 evaluations for each combination of the training dataset, external dataset, phenotype, training sample size, and external sample size, which totaled to over 60 million model evaluations.

### S3. Evaluation in additional datasets

**Figure S2.**
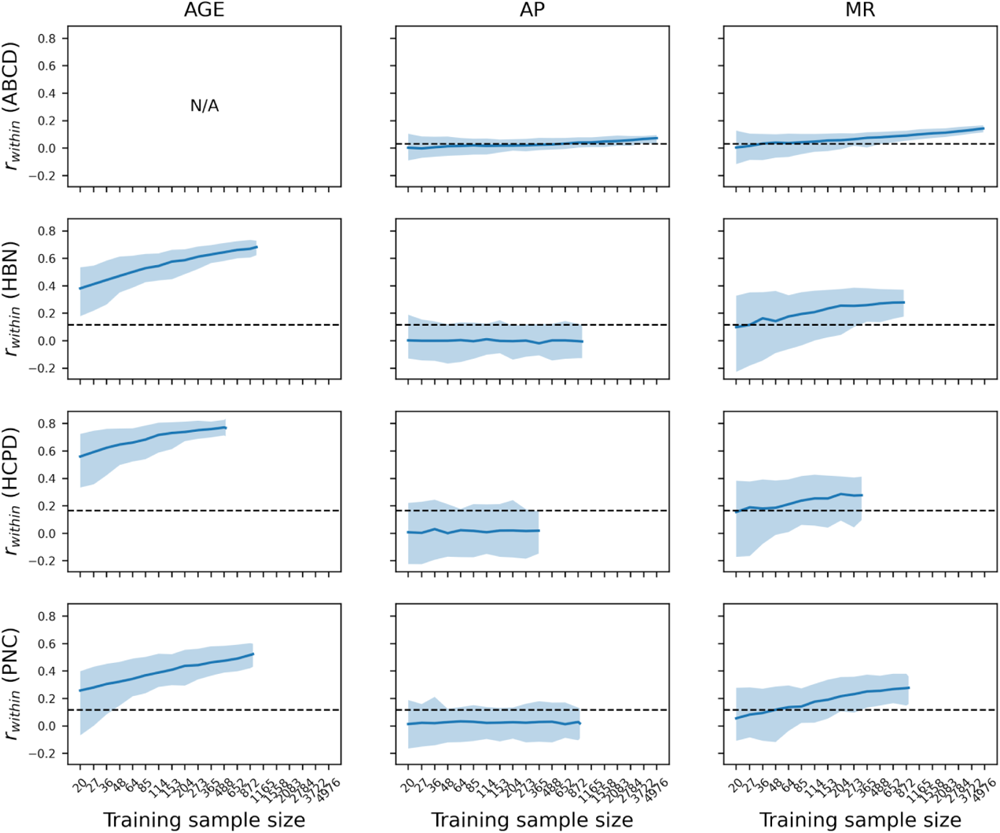
related to Figure 1. Within-dataset held-out prediction performance in all datasets. The performance was evaluated in a randomly selected held-out sample of size n=3000 in ABCD, n=100 in HCPD, and n=200 in PNC. The error bars show the 2.5^th^ and 97.5^th^ percentiles among 100 repeats of resampling at each training sample size. The dotted line reflects the correlation value required for a significance level of *p*<0.05. AP: attention problems, MR: matrix reasoning.

**Figure S3.**
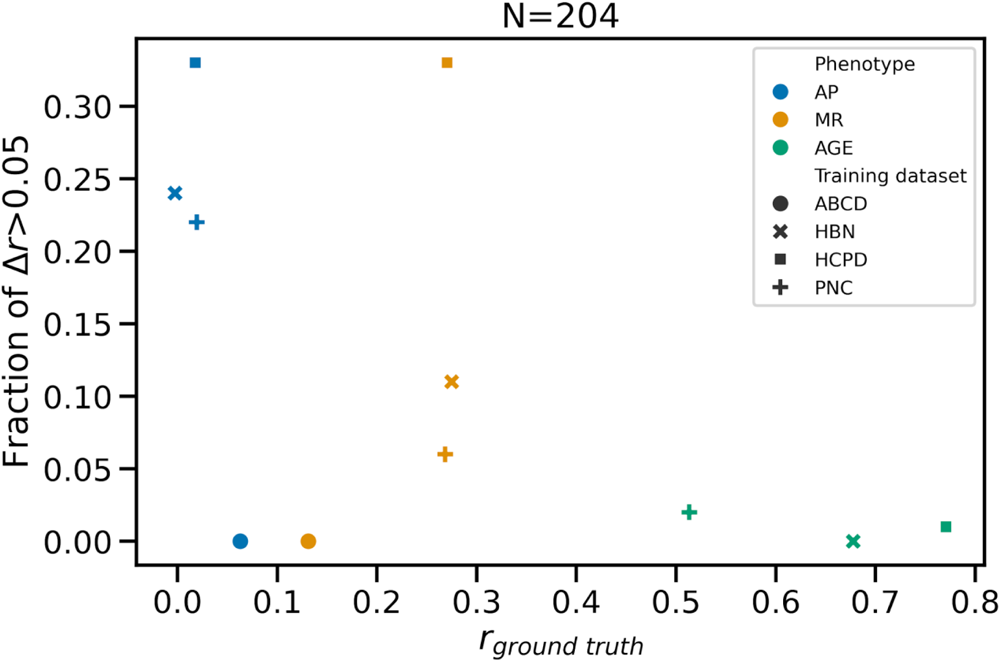
related to Figure 1. Fraction of within-dataset prediction performance exceeding the ground truth by *Δr*>0.05 at a sample size of n=204. AP: attention problems, MR: matrix reasoning.

**Figure S4.**
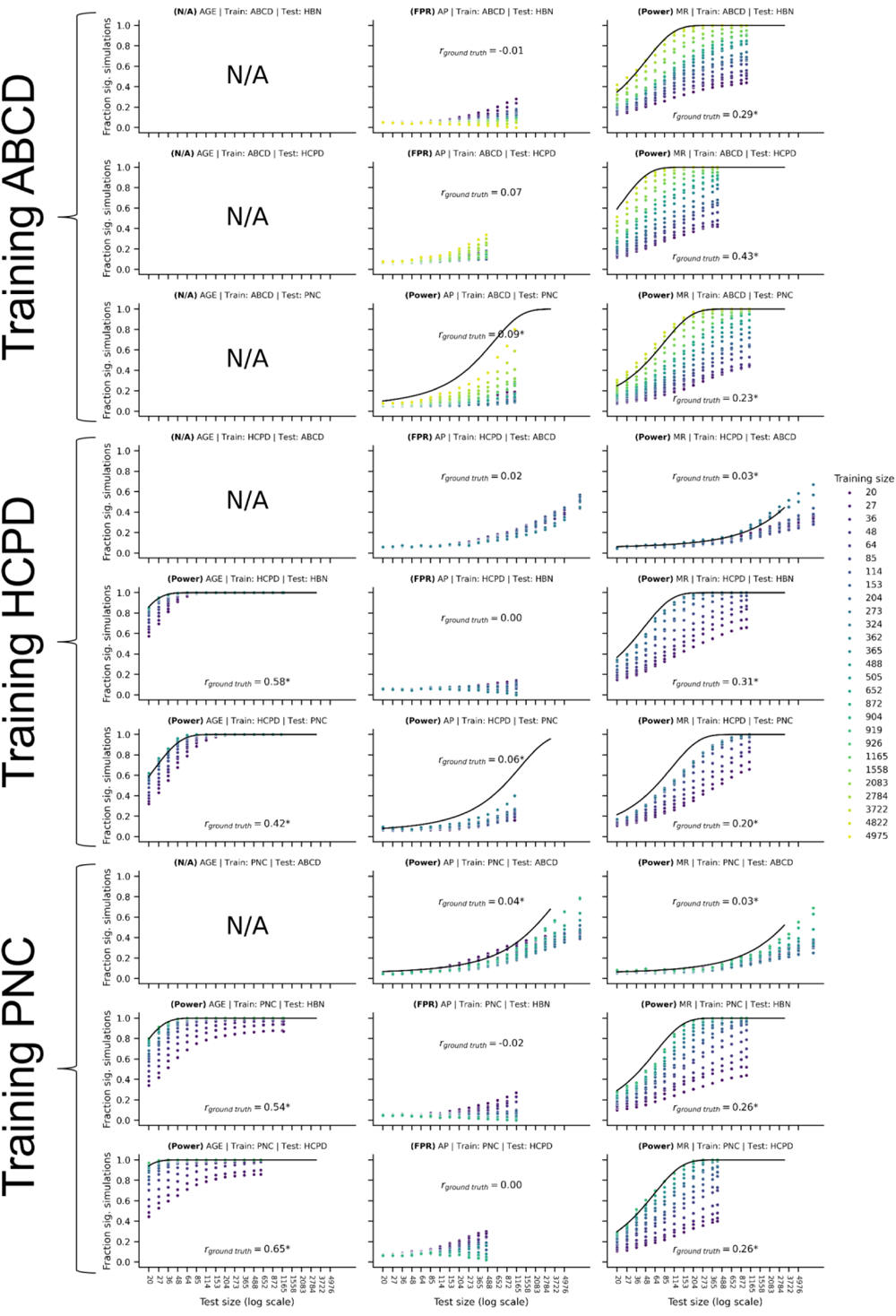
related to Figure 2. Analysis of power and false positive rates when training models in the additional three datasets: ABCD, HCPD, and PNC. Panels with N/A mean that data were not included in this study. AP: attention problems, MR: matrix reasoning.

**Figure S5.**
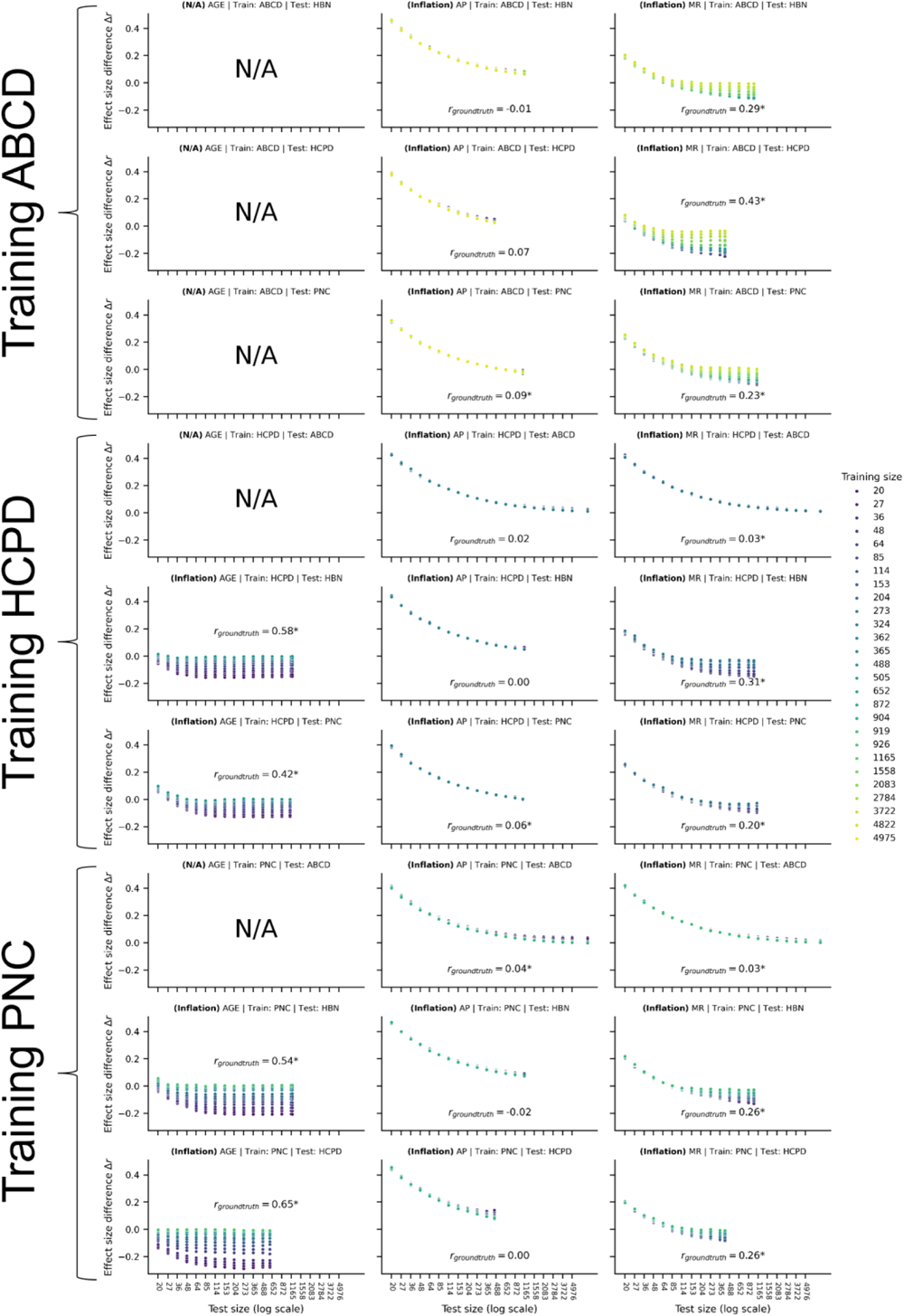
related to Figure 3. Median effect size inflation when training models in the additional three datasets: ABCD, HCPD, and PNC. Panels with N/A mean that data were not available. AP: attention problems, MR: matrix reasoning.

**Figure S6.**
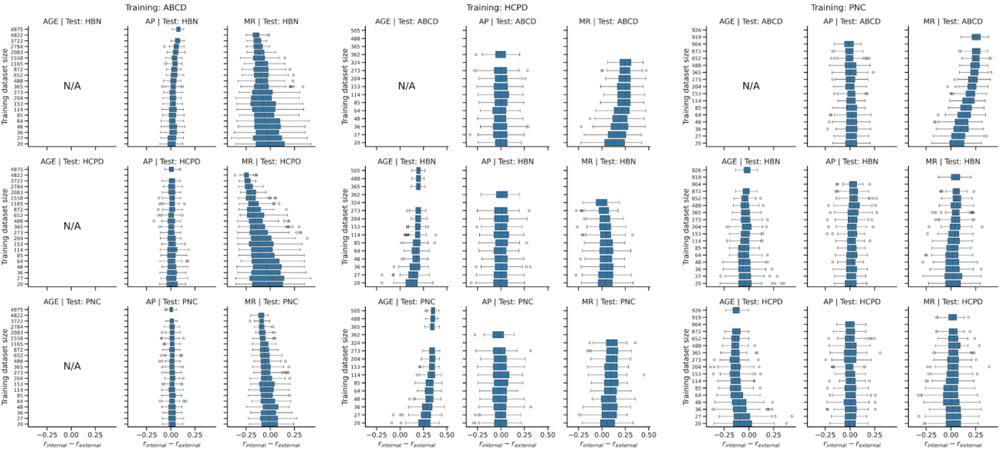
related to Figure 4. Boxplots of the difference between internal and external performance for each subsample of the training data in ABCD, HCPD, and PNC. For each training data size, 100 random subsamples were taken. For internal performance, the model was evaluated in a held-out sample of size n=3000 for ABCD, n=100 for HCPD, and n=200 for PNC. For external performance, the model formed in the training subsample was applied to the full external dataset. Panels with N/A mean that data were not available. AP: attention problems, MR: matrix reasoning.

## S4. Scaled matrix reasoning

**Table S2.** External validation performance in ABCD, HBN, and HCPD for Matrix Reasoning Total Raw Score and Matrix Reasoning Scaled Score. Scaled scores were not available in PNC. *p<0.05, **p<1e-5.

|  | Training Data |  |  |  |  |  |
| --- | --- | --- | --- | --- | --- | --- |
|  | ABCD |  | HBN |  | HCPD |  |
| External Data | MR | MR (scaled) | MR | MR (scaled) | MR | MR (scaled) |
| ABCD | Within | Within | -0.03 | 0.07** | 0.03* | 0.09** |
| HBN | 0.29** | 0.08* | Within | Within | 0.31** | 0.11* |
| HCPD | 0.43** | 0.23** | 0.25** | 0.23** | Within | Within |

**Figure S7.**
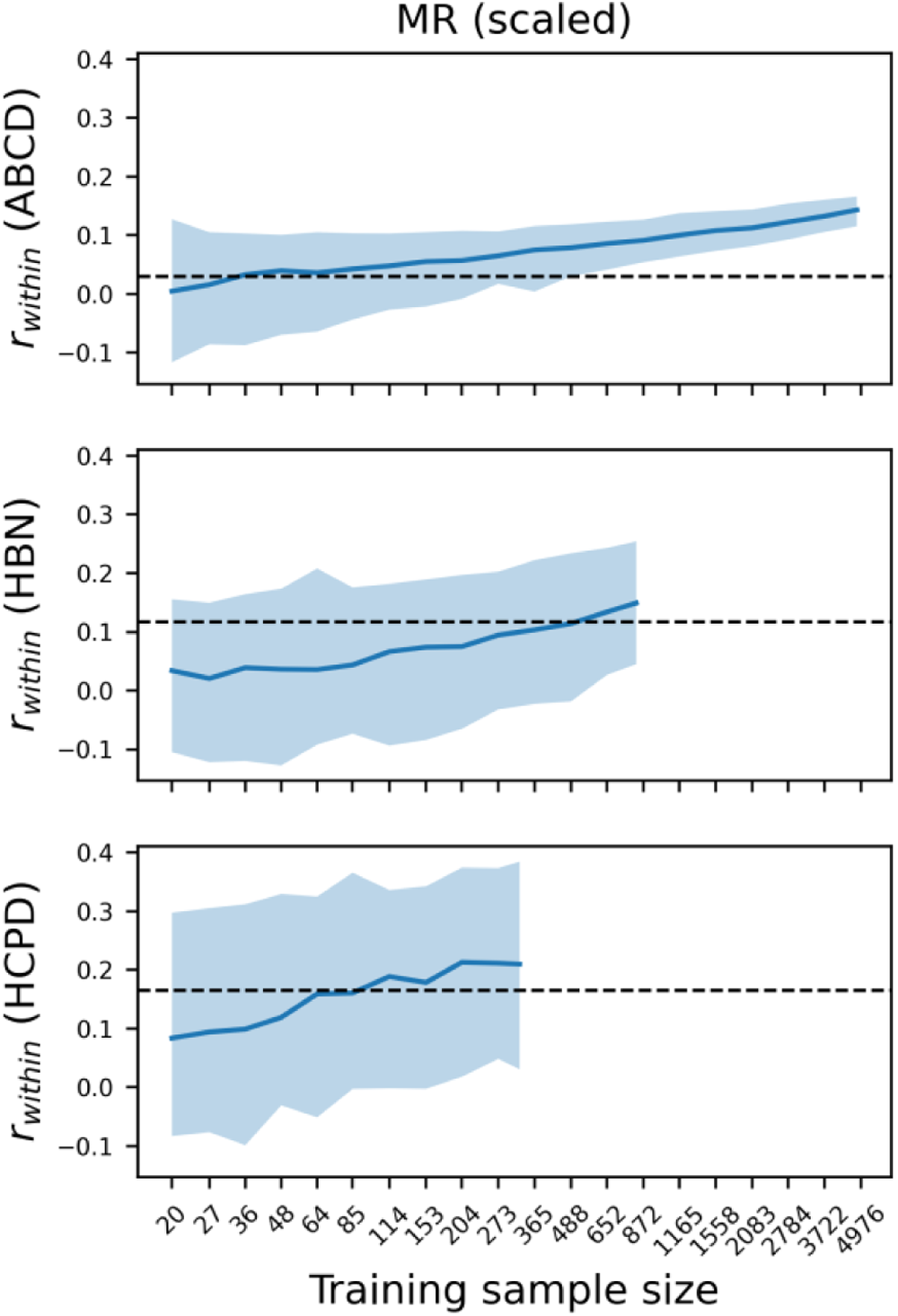
related to Figures 1 and S2. Within-dataset held-out prediction performance in ABCD, HBN, and HCPD for scaled matrix reasoning. In the main text, the total raw matrix reasoning score was used, but here we re-analyzed the data using the scaled score.

**Figure S8.**
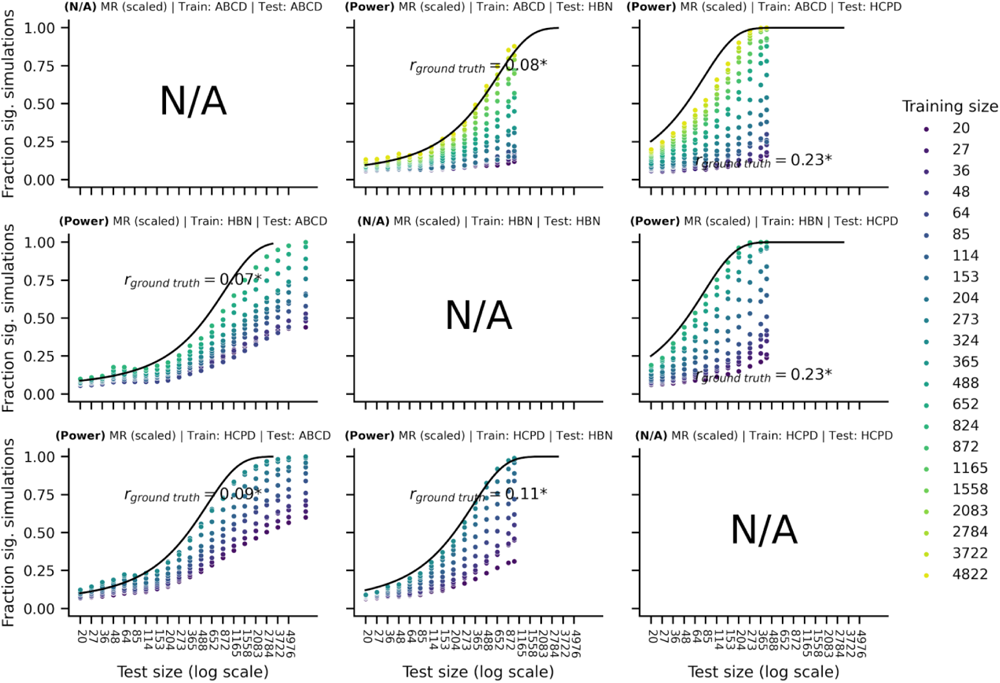
related to Figures 2 and S4. Power and false positive rates for cross-dataset predictions using scaled matrix reasoning. The row reflects the training dataset (ABCD, HBN, HCPD), and the column reflects the test dataset (ABCD, HBN, HCPD). In the main text, the total raw matrix reasoning score was used, but here we re-analyzed the data using the scaled score.

**Figure S9.**
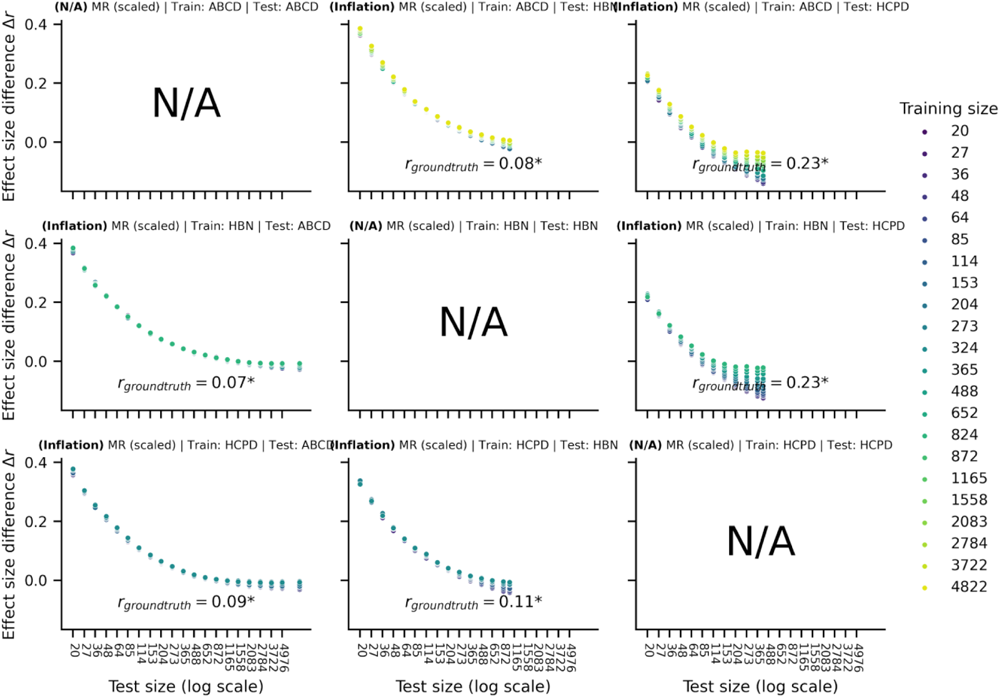
related to Figures 3 and S5. Median effect size inflation for cross-dataset predictions. The row reflects the training dataset (ABCD, HBN, HCPD), and the column reflects the test dataset (ABCD, HBN, HCPD). In the main text, the total raw matrix reasoning score was used, but here we re-analyzed the data using the scaled score.

## S5. Literature review of external validation sample sizes

We performed a brief literature review to contextualize the power and external validation results. Using PubMed, we searched for articles with the following keywords to find functional connectivity prediction papers using external validation: (“functional connect*” OR (“fMRI” AND “connect*”)) AND (“predict*”) AND (“external” OR “cross-dataset” OR “across datasets” OR “generaliz*”). In cases where the articles used multiple training or external datasets, we recorded the sample size of the largest one. Articles were restricted to 2022 and 2023, which returned 117 articles as of July 2023. Articles were excluded for lacking external validation, not using fMRI connectivity data, or inadequate reporting details. Ultimately, 27 articles were included in our sample. The median sample size of the training dataset was n=161 (IQR: 100-495), and the median sample size of the external dataset was n=94 (IQR: 39.5-682). An additional analysis by Yeung et al. included papers before 2022 (Yeung *et al*., 2022), and they found 27 articles using external validation. In this sample, the median sample size of the training dataset was n=87 (IQR: 25-343), and the median sample size of the external dataset was n=137 (IQR: 60-197). In both our dataset and the Yeung et al. dataset combined, the median training sample size was n=129 (IQR: 59.5-371.25), and the median external sample size was n=108 (IQR: 50-281).

